## Supplementary figures and images for "Aggregation of HAPLN2, a component of the perinodal extracellular matrix, is a hallmark of physiological brain aging in mice"

### S1 Fig

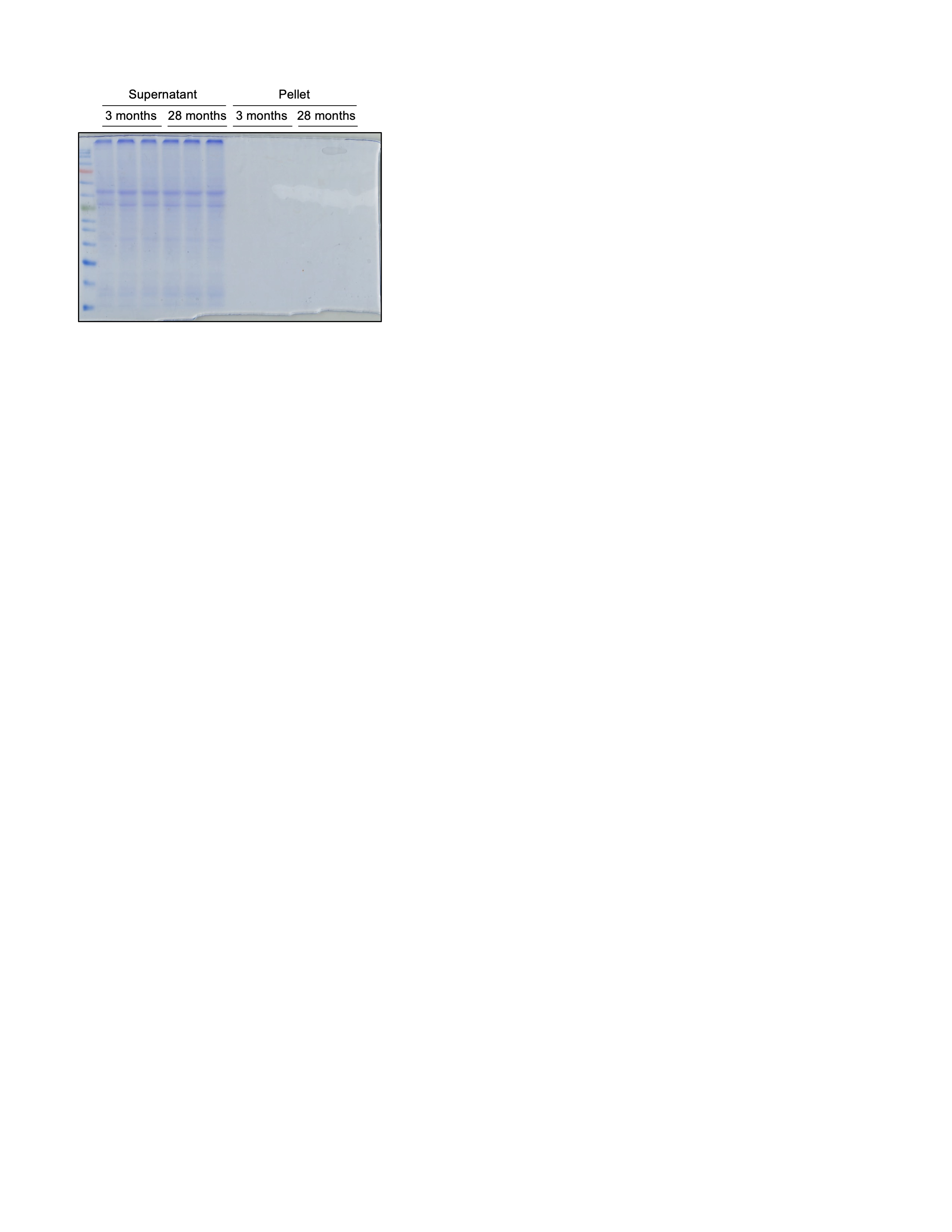

### S2A Fig

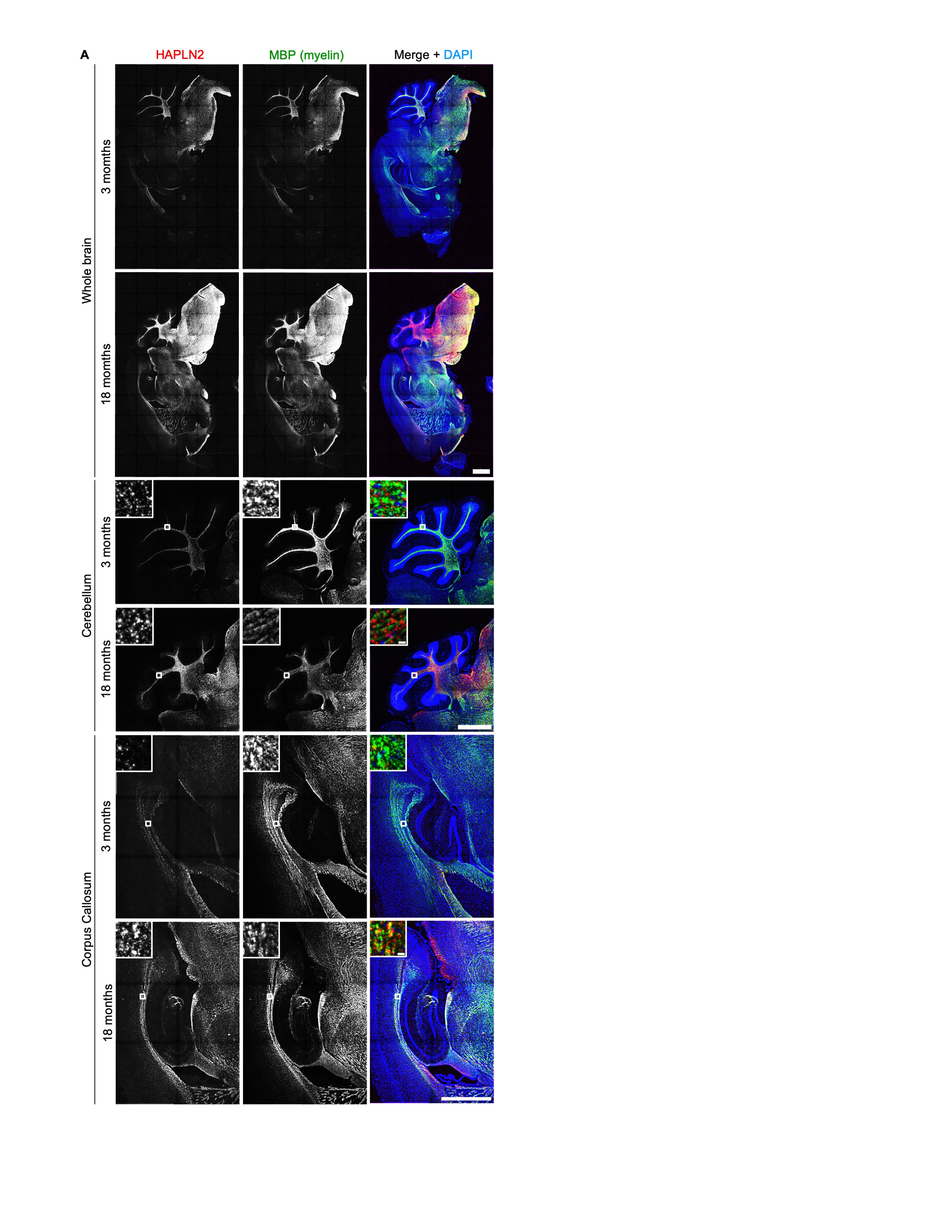

### S2B-D Fig

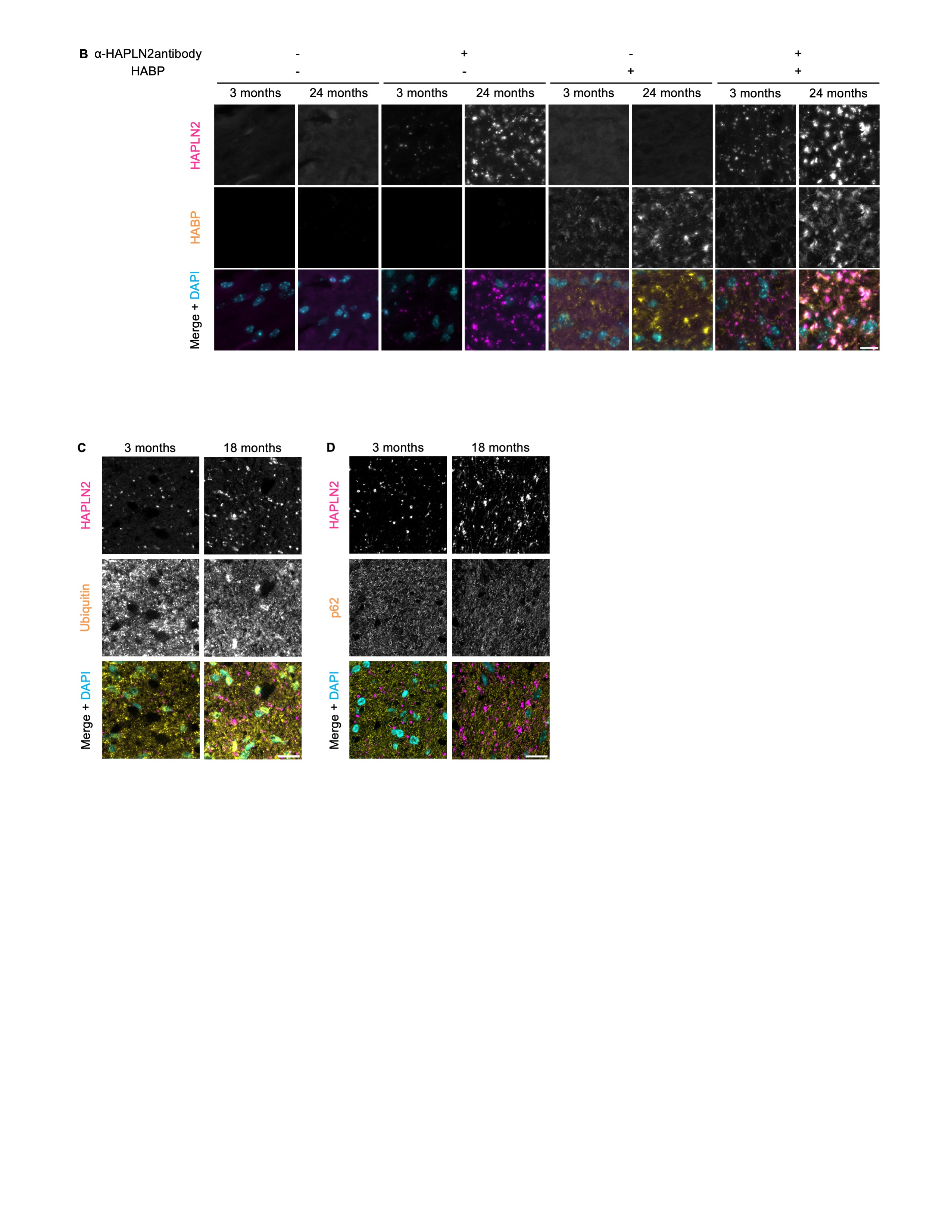

### S3 Fig

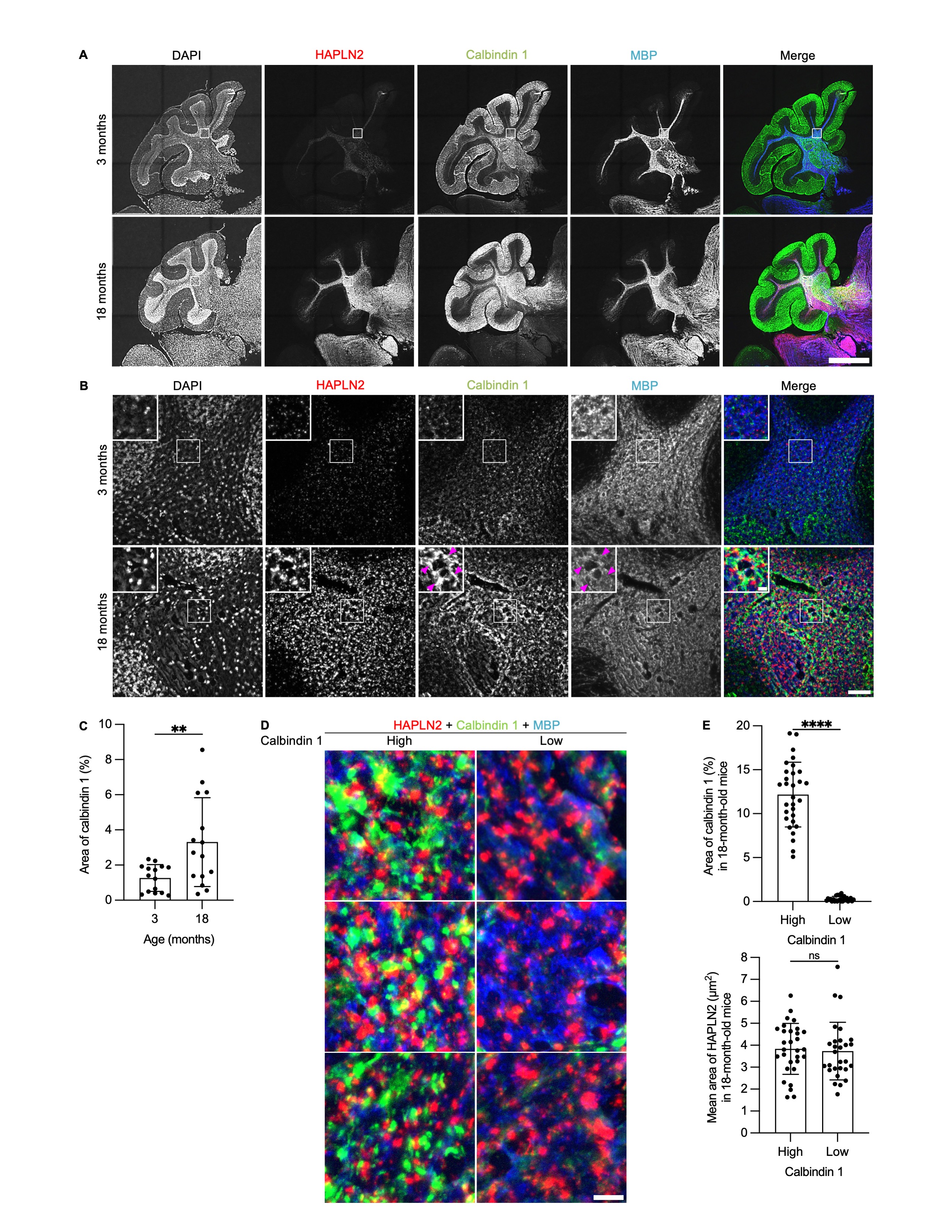

### S4 Fig

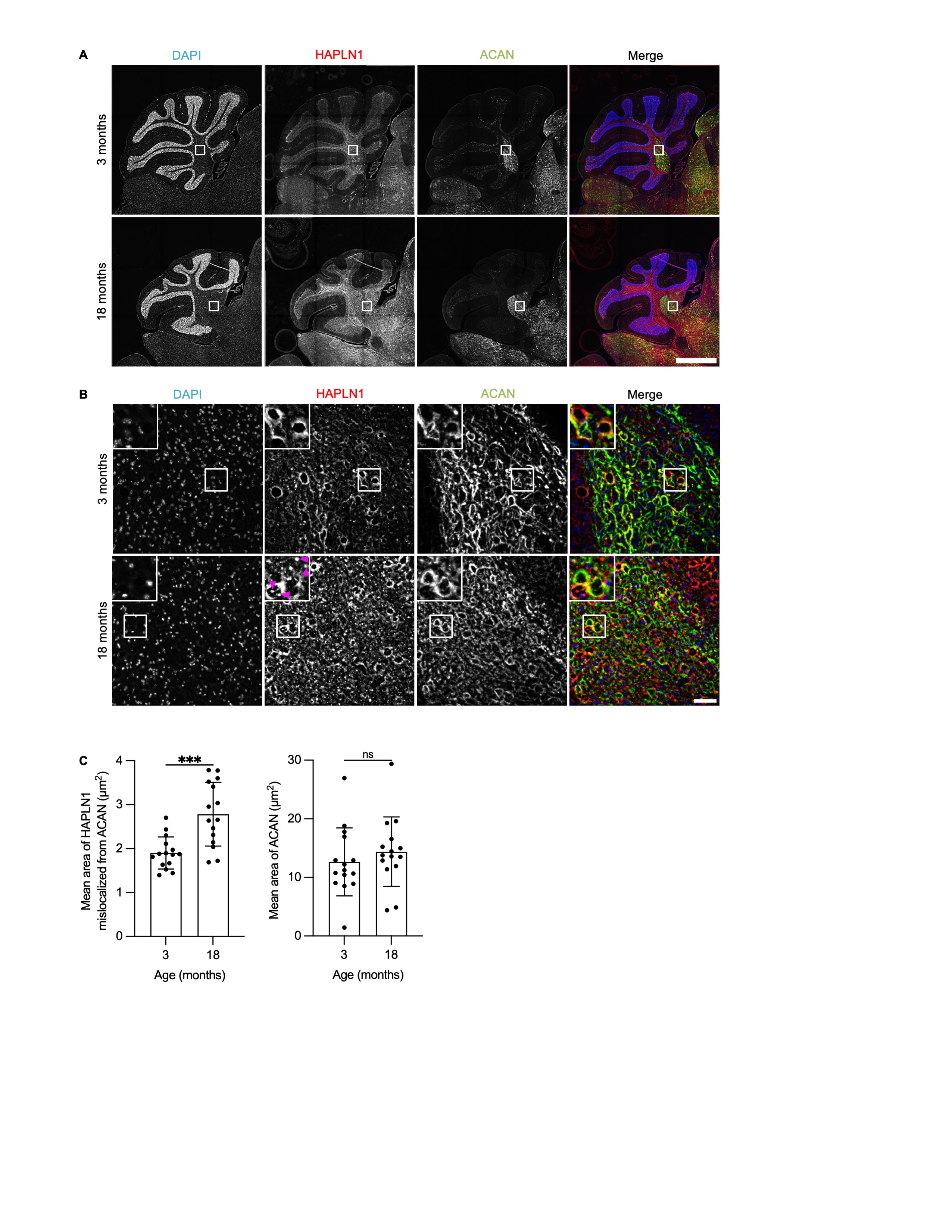

### S5 Fig

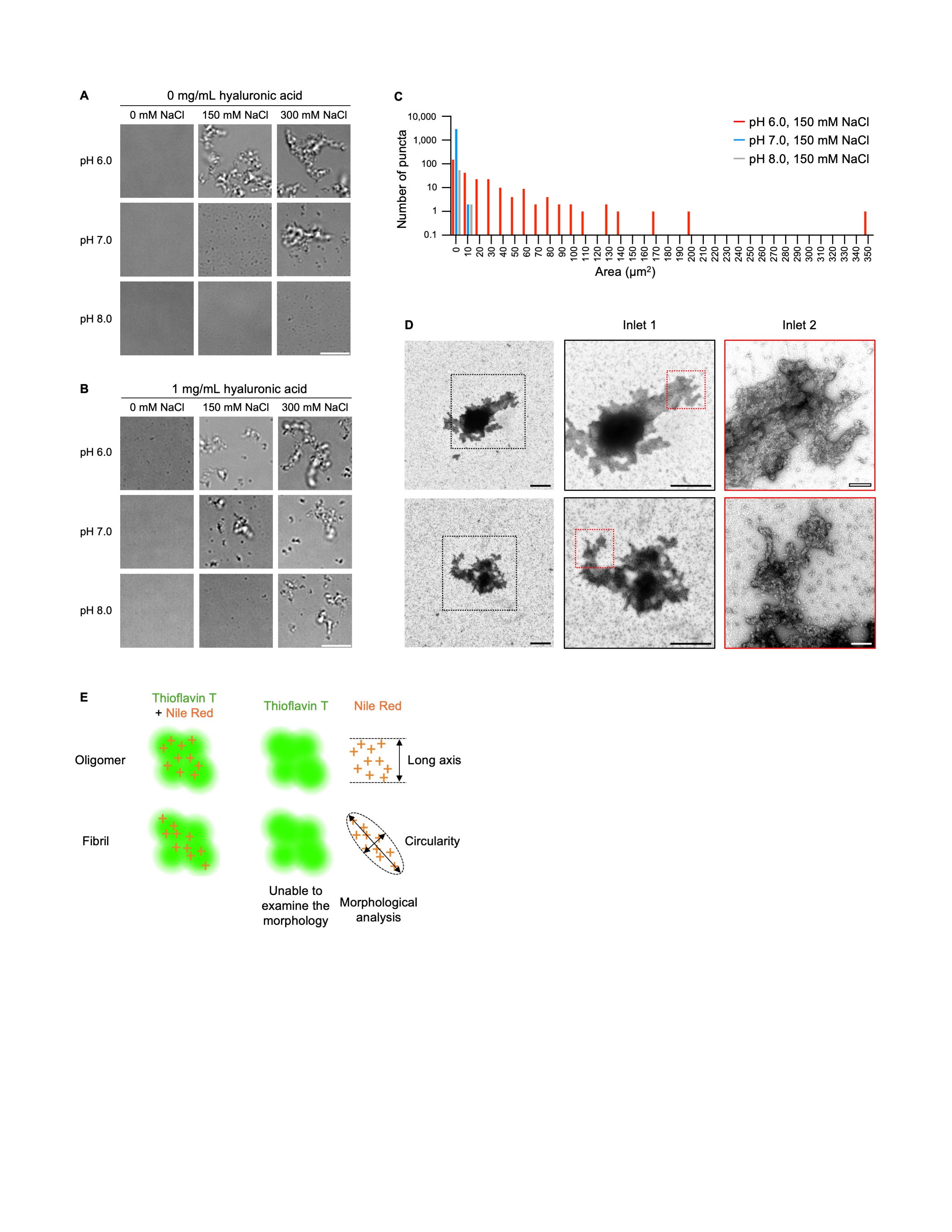

### S6 Fig

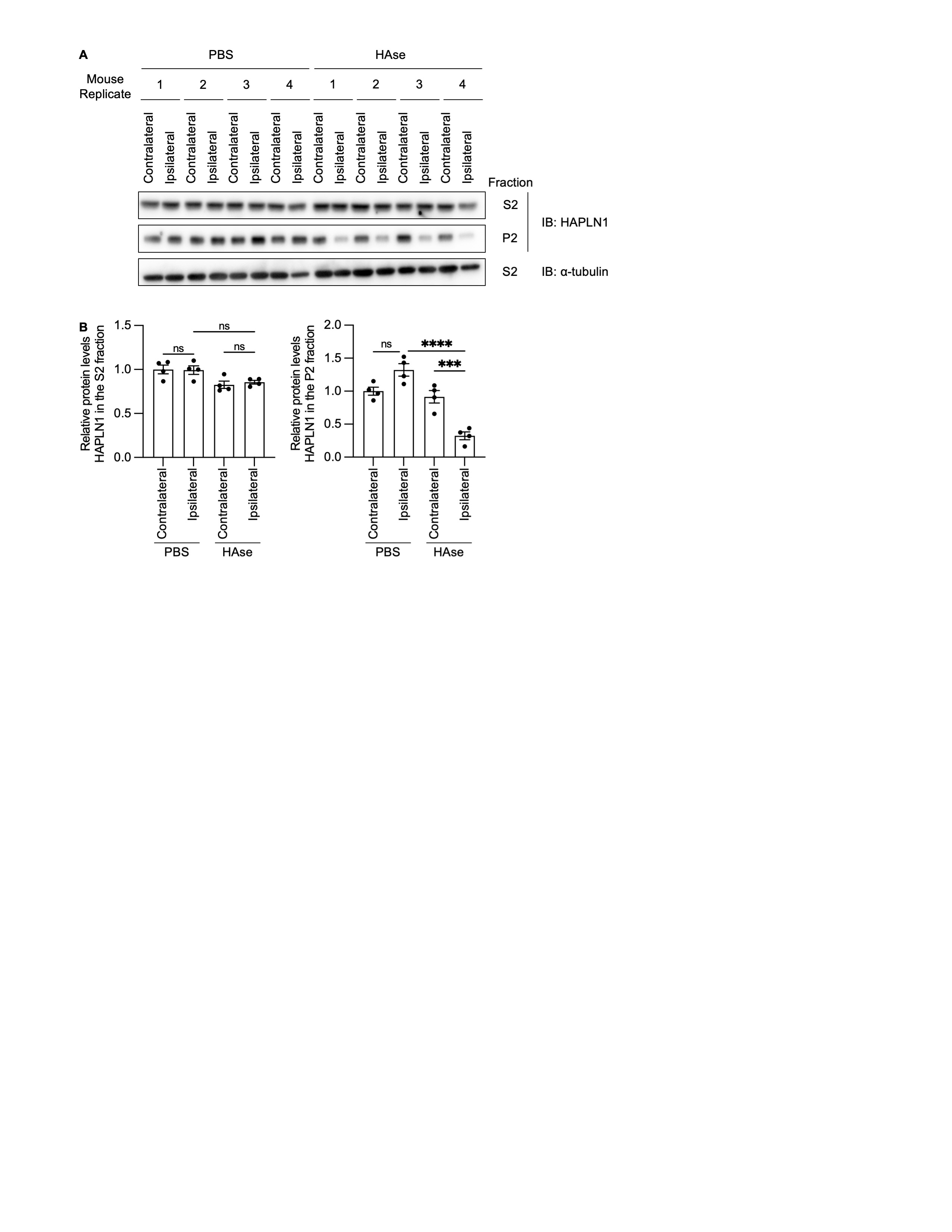

### S7 Fig

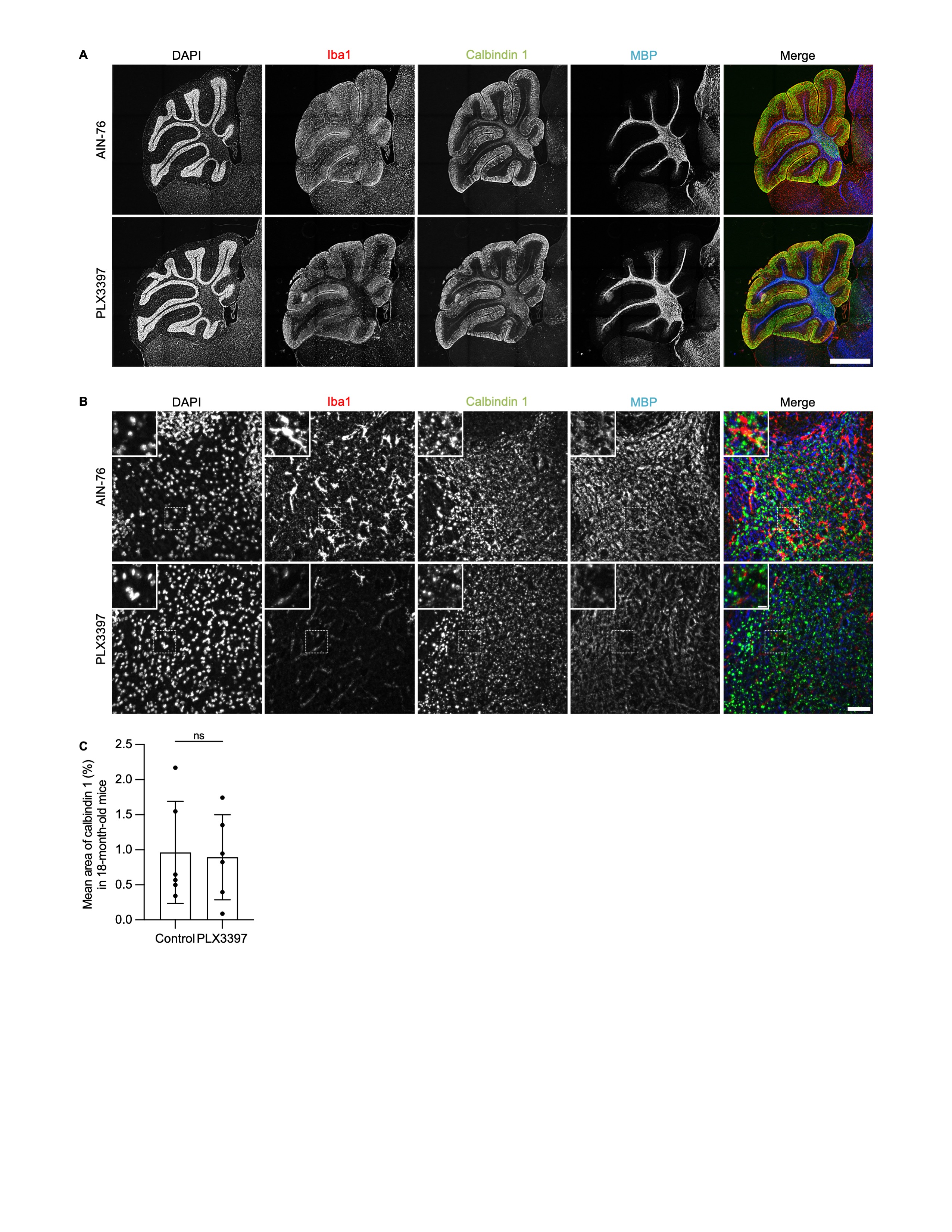

### S8 Fig

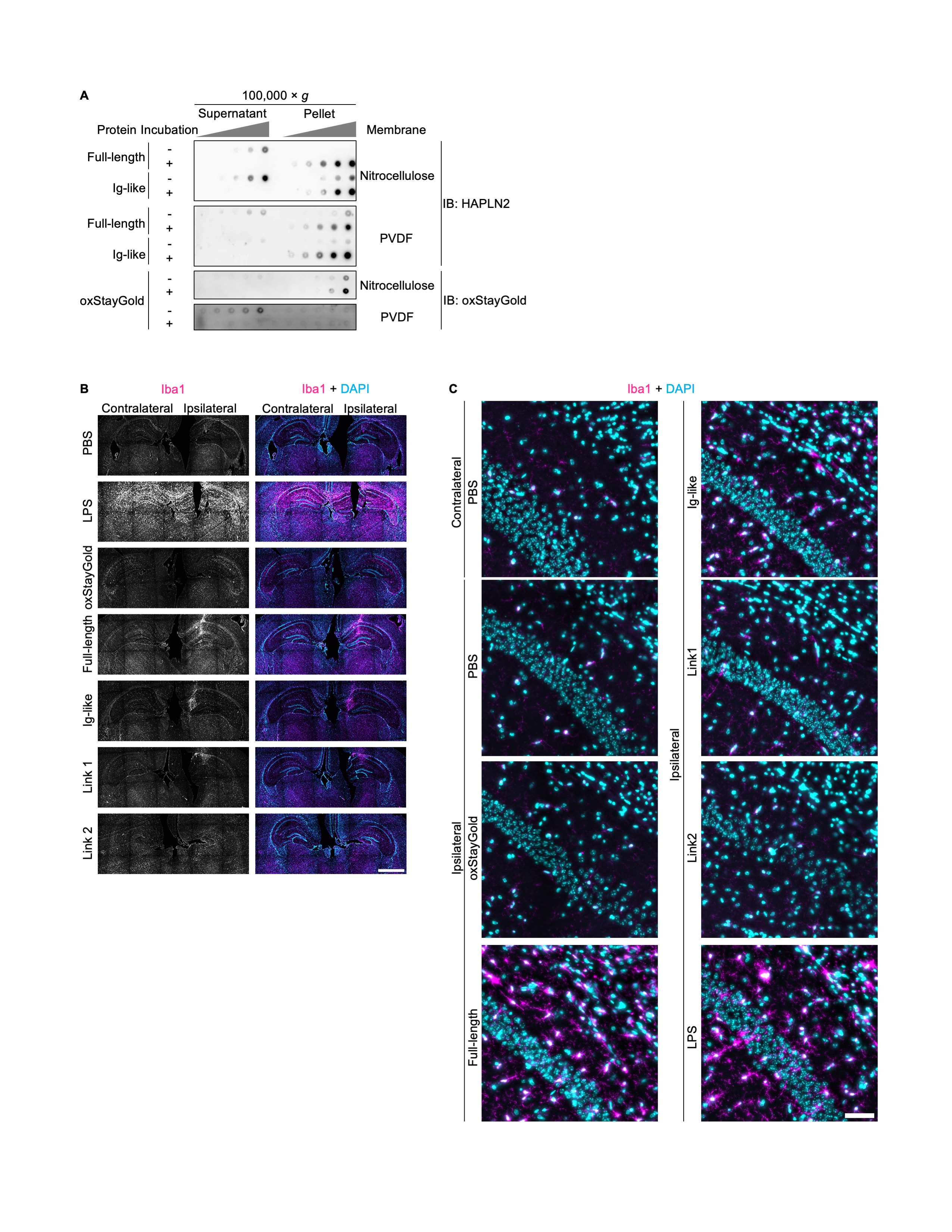
